# Automated behavior classification of *julius seizure* mutants in *Drosophila* reveals stereotyped seizure stages with genotype specificity

**DOI:** 10.1101/2025.05.13.653908

**Authors:** Huiqi Zhang, David L. Deitcher, Chunyue Geoffrey Lau

## Abstract

Bang-sensitive (BS) *Drosophila* mutants exhibit a stereotyped pattern of seizure behavior after mechanical disturbances. We previously identified mutations in the *julius seizure* (*jus*) gene, formerly *CG14509*, can induce BS seizures. However, the behavioral manifestations of the seizure phenotype of the various *jus* mutants have not been fully characterized. Here, we developed a machine learning pipeline featuring LASC (<u>L</u>ong short-term memory and <u>A</u>ttention mechanism for <u>S</u>equence <u>C</u>lassification) for automatic phenotyping of *jus* mutant videos. LASC achieves 90% classification accuracy in distinguishing five phases: paralysis (P), tonic seizure (T), spasm (S), clonic seizure (C), and recovery (R). Applying the trained LASC model to multiple *jus* lines showed they use a common repertoire of seizure stages and followed the general P→T→S→C→R progression, but each genotype exhibited unique patterns of stage duration and transition probabilities. Remarkably, stage usage patterns are distinct among the mutant genotypes. These findings establish that while all *jus* mutants adhere to stereotyped behavioral rules, each allele generates a distinct signature in stage usage. This work demonstrates how advanced behavioral quantification can reveal previously hidden relationships between gene mutation and complex motor outputs. More broadly, the complete pipeline presented here can pave the way for high-throughput, automated drug screening for epilepsy.

## Introduction

Epilepsy affects approximately 46 million people worldwide across all age groups, making it one of the most common neurological diseases globally ^1^. Although drugs are available for treating epilepsy, about one-third of patients are resistant to current anti-epileptic drugs. To elucidate the mechanisms of epilepsy and develop novel therapies, researchers widely employ animal models ^2,3^. The fruit fly *Drosophila melanogaster* is an attractive model system due to its short generation time, relatively low cost, and ease of genetic manipulation. Over the years, several seizure-associated gene mutations have been discovered in *Drosophila* and utilized in epilepsy research ^4–9^. These flies exhibit remarkable seizure behavior and a refractory period after mechanical shock. A brief, strong mechanical shock – referred to as a “Bang” – can be induced by vigorously tapping or vortex-mixing the fly vial. Such Bang-sensitive (BS) seizures resemble rodent models of electric kindling or pharmacologically induced seizures, as well as drug-resistant epilepsy in humans, underscoring its importance in epilepsy studies ^10^.

Previously, we identified a new mutant in the gene, *julius seizure* (*jus*; formerly CG14509), which confers BS seizure susceptibility ^11,12^. *jus* mutants and other BS mutants both show distinguishable stages within a seizure session. Early studies described these stages as initial seizure, temporary paralysis and recovery seizure ^5^. Later, six phases were identified in the *para*^*bss1*^ mutants: (1) initial seizure; (2) paralysis; (3) tonic-clonic-like activity; (4) recovery seizure; (5) refractory period; (6) complete recovery ^10^. Although these stages were qualitatively well-defined, seizures were traditionally quantified using a single parameter: the time from Bang to complete recovery. Recent advances in deep learning (DL) technology now enable automated, high-resolution tracking of body parts and more refined behavioral quantification ^13–16^.

In this study, we aim to precisely quantify seizure stages and map the relationship between *jus* genotypes and behavioral phenotypes. A key advantage of BS seizures is that they can be immediately induced in a non-invasive manner. Moreover, BS seizure behavior closely mirrors the seizure-like electrophysiological activity in fly mutants ^8,12,17^, validating behavioral analysis as a reliable approach. However, quantifying seizure behaviors in *Drosophila* is challenging due to their small size, rapid movements, and highly variable leg postures. Here, we developed a comprehensive experimental and analytical pipeline, encompassing seizure induction, video acquisition, behavioral quantification, and group comparison. A critical component of this pipeline is a novel seizure stage classification system. To facilitate seizure stage identification, we developed LASC (LSTM and Attention mechanism for Sequence Classification) classifier, a method based on the long short-term memory (LSTM) algorithm. The well-trained LASC model effectively captured stage-specific behavioral patterns and achieved accurate classification. Applying LASC, we generated behavioral sequences for multiple *jus* mutants and compared their stage transitions and behavioral structures.

## Results

### Leg movement patterns characterize seizure stages in *jus* mutants

Here, we examined four different *jus* mutant lines. They were: (1) *PBac*/*Df*, (2) *PBac*, (3) *jus* ^*iso7*.*8*^/*Df*, and (4) *jus* ^*iso7*.*8*^/ *PBac*. The *PBac* allele (fully termed *PBac[WH]CG14509*^*04904*^) carries a transposon insertion in *jus* and acts as a strong hypomorph. The *PBac* line is homozygous for this insertion. The *Df* allele (formally named *Df(3R)BSC501*) deletes the entire *jus* locus, making *PBac/Df* a hemizygous combination of the *PBac* insertion and the deletion. *jus*^*iso7*.*8*^ is a weak hypomorph, so *jus*^*iso7*.*8*^*/Df* and *jus*^*iso7*.*8*^*/Pbac* represent allelic combinations with the deletion or *PBac* insertion allele, respectively.

Previously, we reported distinct recovery times across these lines: *PBac/Df* > *PBac* > *jus*^*iso7*.*8*^*/Df* > *jus*^*iso7*.*8*^*/PBac* ^12^. Based on this, we hypothesized that the four strains would exhibit distinct behavioral phenotypes. To investigate motor behavior, we prepared flies, induced seizures and recorded high-speed videos (Fig. 1 and Methods). Flies residing in the vials were subjected to 10 sec of vortex mixing, immobilized dorsal side down, and seizures were recorded from the ventral side at a frame rate of 100 Hz to capture fast leg movements.

**Figure 1.**
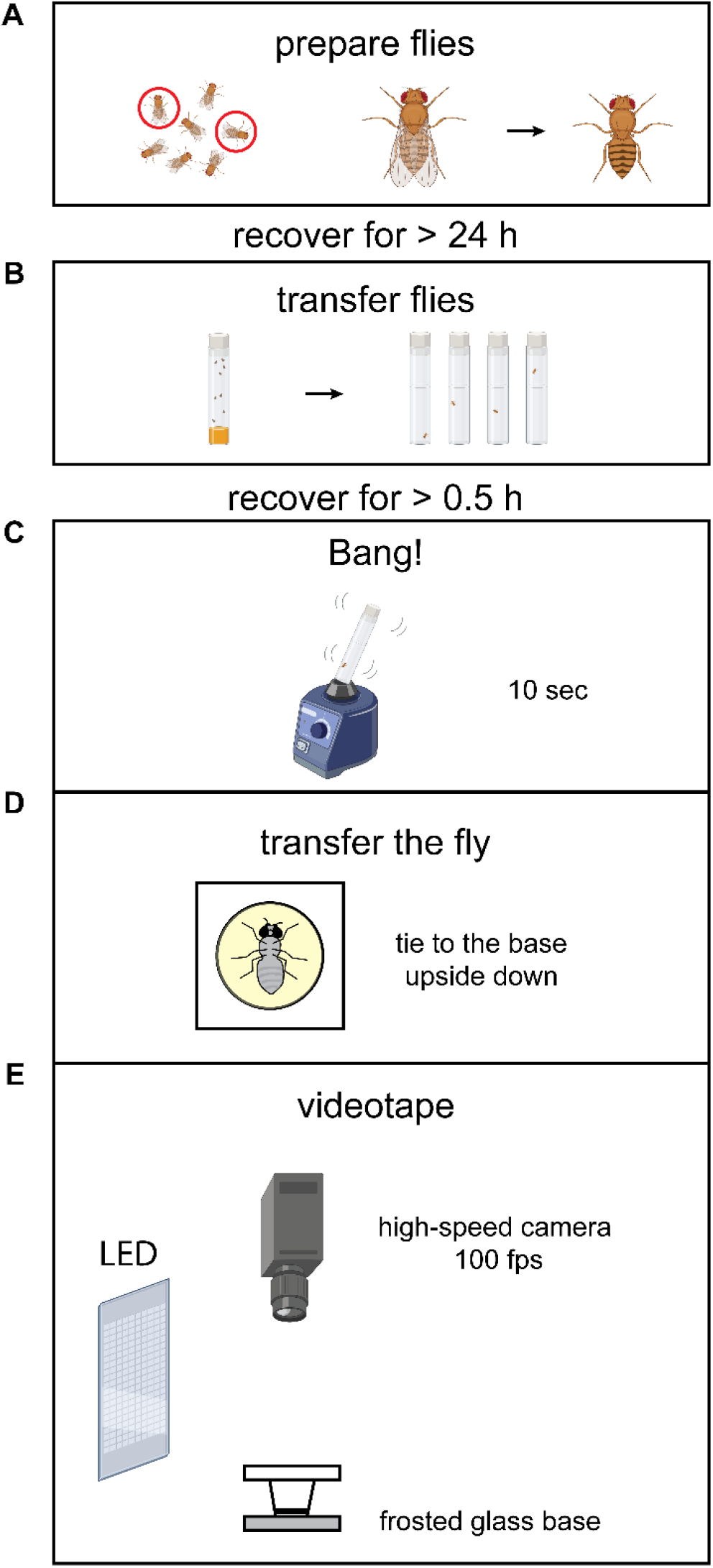
Subject preparation, seizure induction and video acquisition. (**A**) *jus* mutants were anesthetized under CO_2_, selected and their wings were clipped. (**B**) After 24 hours recovery, flies were isolated in empty vials and followed by 0.5 h recovery. (**C**) 10-second vortex mixing was applied to induce seizures. (**D**) Flies were transferred to the recording chamber and tethered to the base upside down. (**E**) Recording apparatus. Seizures were recorded with a high-speed camera (100 frames per second).

Seizures in all four genotypes were manifested through evident leg movements. When a 10-second vortexing stimulus was used, the initial seizure was not observed, because the initial responses were finished before vortexing ends, which were also reported in *para*^*bss1*^ and *eas* mutants ^8,18^. Based on visual inspection of body movement, we defined five stages in between vortexing end and full recovery (Table 1). For example, in a *PBac* mutant, paralysis (P, Video 1) was closely followed by tonic seizures (T, Video 2), and later with an extended duration of spasms (S, Video 3), clonic seizures (C, Video 4) and recovery episodes (R, Video 5). Similar to *para*^*bss1*^ mutants, the *jus* mutants displayed spasm and tonic-clonic seizures. They also shared traits with other BS mutants (*para*^*bss1*^, *eas*, and *tko*^*25t*^), including rolled-up abdomen, proboscis extensions and females laying eggs ^8,18,19^. However, unlike these mutants, which show fast wing flapping and scissoring, *jus* seizures did not involve wing movements. Therefore, the wings were removed during preparation for better contrast in the videos. Drawing from these observations, we focused on the leg dynamics in our analysis.

**Table 1.** Seizure stages in *jus* mutants.

| Stage | Definition |
| --- | --- |
| Paralysis (P) | Immobility of all body parts |
| Tonic seizure (T) | Stiff legs with uncoordinated, non-rhythmic and large leg movements |
| Spasm (S) | Rhythmic or non-rhythmic small random leg movements |
| Clonic seizure (C) | Rhythmic, synchronous small leg movements |
| Recovery episode (R) | Normal, continuous leg movements preceding either additional seizure stages or full recovery |

We used Deeplabcut (DLC) to track fly’s body parts^20^. To fully extract a fly’s movement dynamics, we tracked 31 body parts, including their head, eyes, neck, thorax, abdomen, and leg joints and tips (Supplementary Fig. 1). A DLC model was trained and used to estimate the X and Y coordinates of each selected body part in the video frame by frame. Representative tracking of leg tips in a *PBac* fly (Fig. 2A) revealed complex dynamics on each individual leg. Zoomed-in traces highlighted unique patterns and scales during tonic, spasm, clonic, and recovery phases (Fig. 2B), confirming that leg movements are reflective of seizure stages.

**Figure 2.**
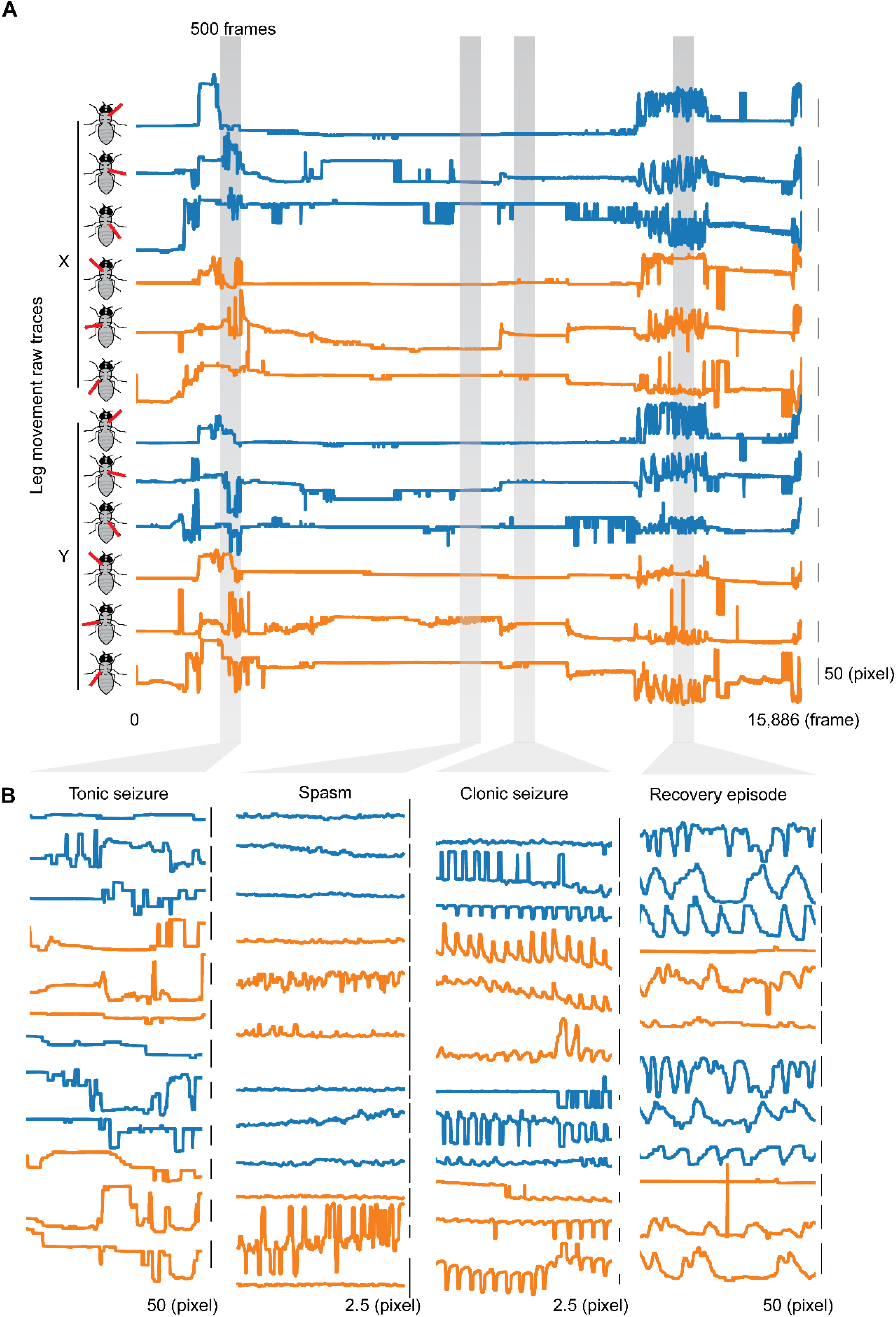
Representative leg movement of *jus* seizure. (**A**) Representative tracking data showing 6 leg tips (both X and Y coordinates) over an entire seizure from a *PBac* fly (total of 15,886 frames or 158.86 sec). Blue and orange traces were the left and right legs, respectively. Plots were in slightly different scales to show leg dynamics properly. (**B**) Magnification of 4 stages in a *jus* seizure, which were tonic seizure, spasm, clonic seizure and recovery episode. 500-frame (5-sec) data of each stage were magnified. Plots were in different scales, with leg movement of each stage clearly visible no matter the scale of movement. The small leg movements, such as those in spasms and clonic seizures, were reliably tracked.

### Behavior analysis pipeline: development of the LASC model

To automatically classify seizure stages from leg movements, we performed time-series classification on DLC-tracked data. While both supervised and unsupervised machine learning algorithms are viable for behavioral time-series analysis ^21–27^, we opted for a supervised approach since BS seizures are not natural behaviors and we had clear expectations about the stages involved. To this end, we developed LASC (<u>L</u>ong short-term memory and <u>A</u>ttention mechanism for <u>S</u>equence <u>C</u>lassification), a model based on long short-term memory (LSTM). We chose the LSTM model because behavioral sequences containing patterns tend to occur over a finite period and LSTM has shown good performance on capturing these long-term dependencies ^28,29^. The workflow of implementing LASC includes: (a) body part tracking (DLC); extraction of motion features; (c) creation of a manually labeled dataset with predefined actions; (d) LASC model training; (e) and behavioral sequencing (Fig. 3A). Applying LASC on *jus* seizures, we extracted features to capture the comprehensive dynamics of individual and paired leg dynamics. The extracted motion features included: (1) corrected distance; (2) speed; (3) acceleration; (4) angle of the tibia-tarsus joint (TTJ); (5) angular speed of the TTJ; (6) the angle between femur (vector from leg base to femur-tibia joint) and body-center axis (vector from abdomen tip to neck); (7) Pearson correlation between every two leg tips (see Methods). These features exhibited distinct patterns across seizure stages (Fig. 3C).

**Figure 3.**
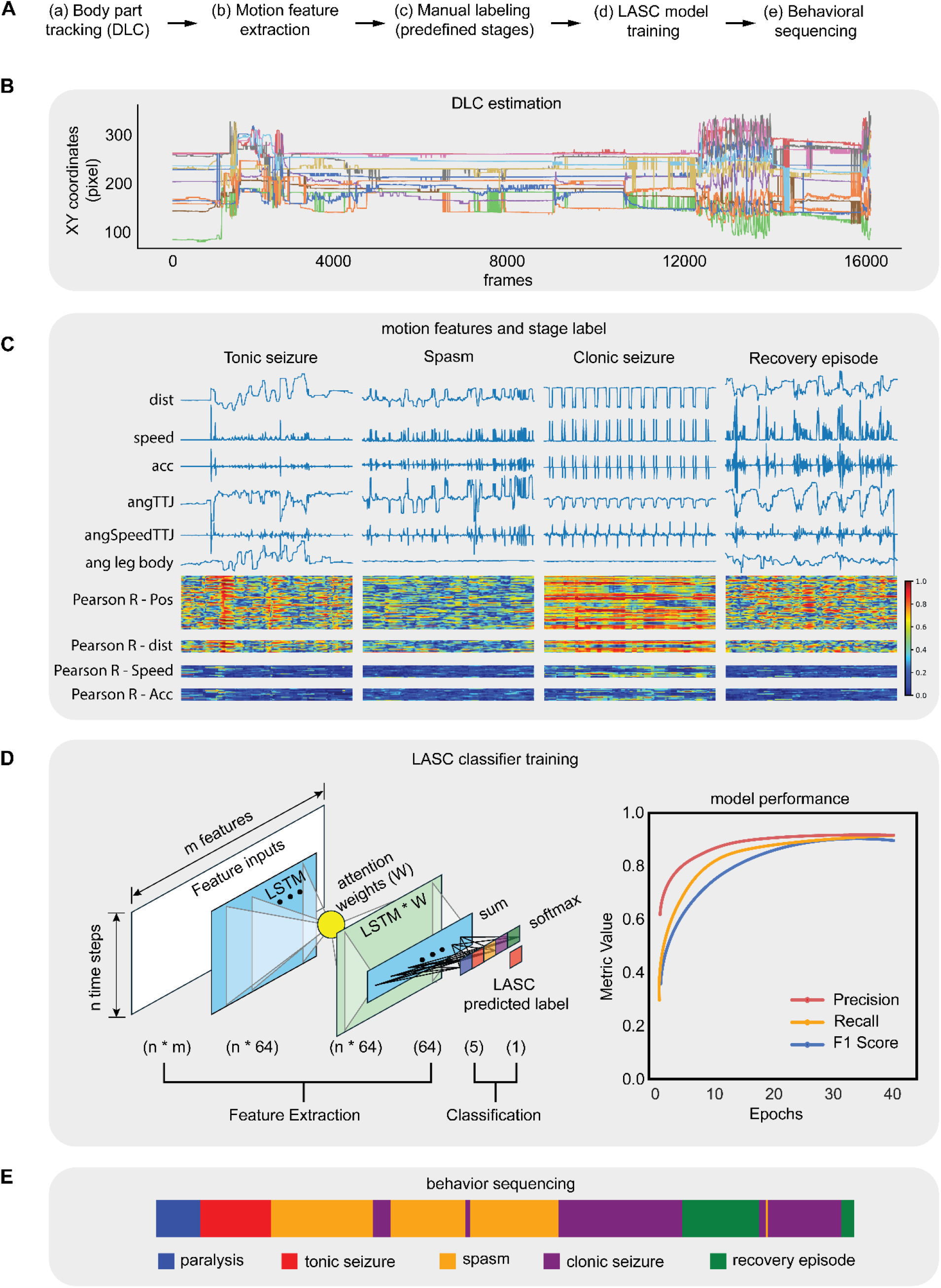
LASC workflow (<u>L</u>ong short-term memory and <u>A</u>ttention mechanism for <u>S</u>equence <u>C</u>lassification). (**A**) LASC is a model that we built to implement behavioral classification after pose estimation. First, the X and Y coordinates of the selected body parts were predicted by DeepLabCut (DLC) from raw videos. Then the tracking data were used to extract motion features, such as speed, acceleration, angles, etc. To perform supervised behavioral classification, behavioral labels were manually generated and concatenate with the motion features. The labelled datasets were fed into the LASC model for training. The trained LASC model was used to implement behavior sequencing. (**B-E**) Applying LASC to sequence *jus* seizure behavior. Here we show the representative outputs of each step. (**B**) Example traces of leg tips (12 traces from X and Y of 6 legs) estimated by DLC. The representative fly is from *PBac* line. (**C**) Representative segments of motion features and corresponding stage labels. The DLC outputs were used to calculate the time series of locomotion, including corrected distance, speed, acceleration (acc), joint angle, angular speed of the tibia-tarsus joint (TTJ), the angle between leg and body, the Pearson correlation (R) of the same type of features between every two legs. Each blue trace was from one leg. Plots were in arbitrary units, with the patterns of the motion features shown clearly. The Pearson correlation heatmaps were from all pairs in a 5-sec time window. The motion features manifested distinct dynamics in each stage. (**D**) Motion features were fed into the LASC model for latent feature extraction and behavioral classification. The left panel showed the layer structure of LASC model. To evaluate the model performance, standard machine learning metrics, precision, recall and F1 score were used. Videos with predicted labels were also generated for visual validation. (**E**) The trained LASC model can be used for sequence an entire video and batch analyses.

Thus, our approach defined each frame as a vector of 159 motion features. The LASC framework first encodes these motion features into deep temporal representations using LSTM, then an attention mechanism is integrated to selectively emphasize informative latent features (Fig. 3D). LASC model performance is quantitatively assessed through conventional classification metrics (precision, recall, weighted F1 score). We also conducted qualitative validation by generating annotated videos that overlayed predicted stage labels onto raw videos. The trained LASC model enabled automated video sequence analysis (Fig. 3E) and high-throughput batch processing of multiple recordings.

### *jus* mutants use a common repository of stages in seizure

Our application of the trained LASC model to 13 flies from four *jus* lines provided a global view of the temporal progression of seizures (Fig. 4A). A majority (8/13) of the flies progressed through all the 5 stages. Flies may experience multiple bouts of clonic seizures and recovery episodes before full recovery. The *jus*^*iso7*.*8*^*/PBac* mutants frequently lacked observable paralysis at the beginning, likely due to the particularly short nature of this phase in the weak hypomorph (paralysis was present but ended before the videotaping started). Some individuals across strains did not show a recovery episode at the end, that were partially due to the individual’s transient recovery episode and partially obscured by smoothing procedure. To the best of our knowledge, we are the first to distinguish between the tonic and clonic sub-phases.

**Figure 4.**
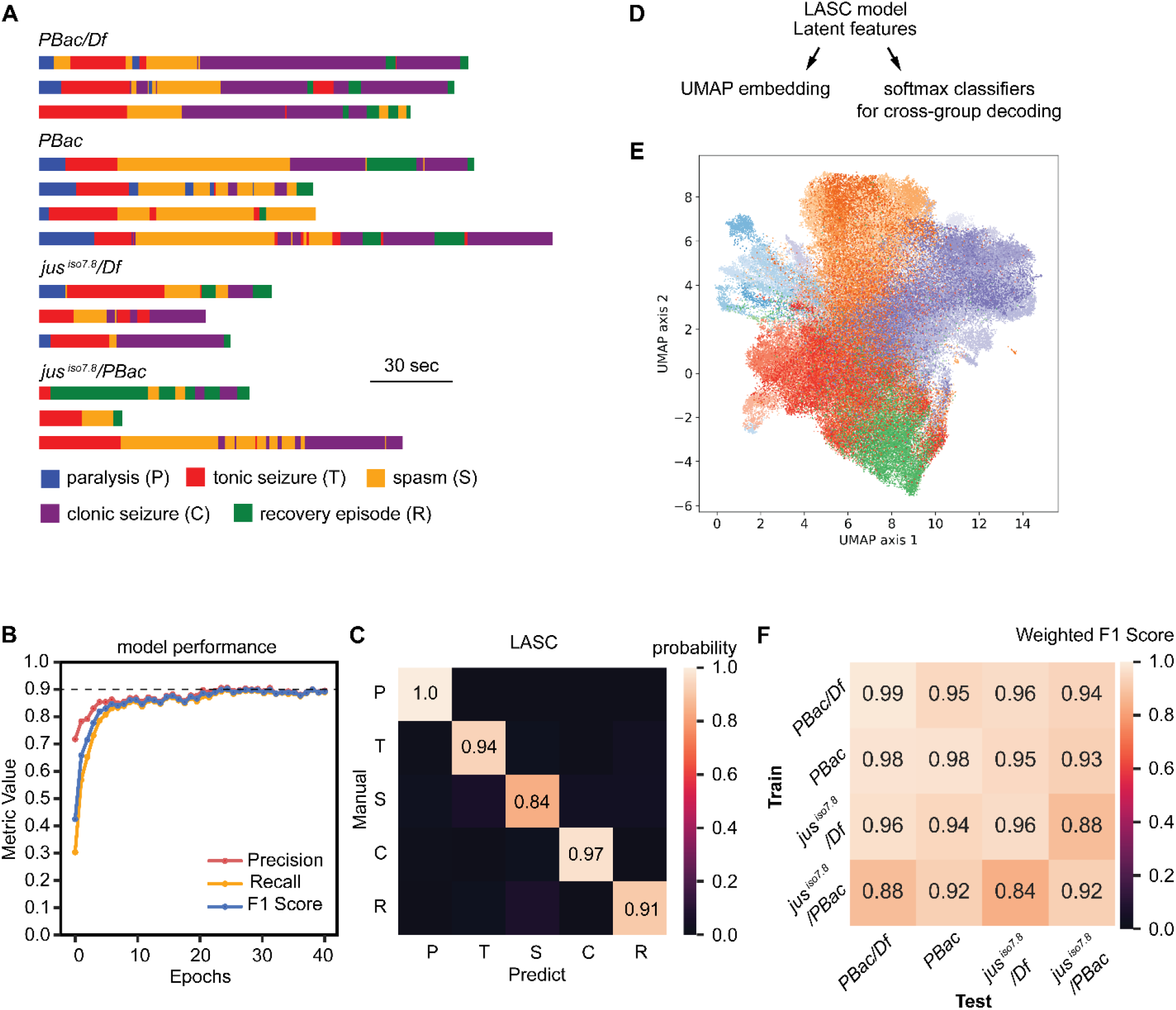
Seizure stages characterized by leg movements are shared across *jus* lines. (**A**) LASC estimated stage sequences of *jus* mutants. Each row represented a fly. Dark blue represents paralysis (P); Red represents tonic seizure (T); Orange represents spasm (S); Purple represents clonic seizure (C); Green represents recovery episode (R). (**B**) Standard machine learning metrics were used to evaluate the LASC classifier performance, including precision, recall and weighted F1 score. (**C**) Confusion matrix between manually labelled ground truth and LASC prediction. The values indicated what fraction of manually labelled data were classified as each stage by the LASC model in the test dataset. (**D**) Pipeline for analyzing seizure stage classification in four *jus* lines by using latent features from the LASC model. UMAP for visualization and cross-group decoding test for quantifying the stage shareability. (**E**) UMAP embedding of the latent features from 4 lines (same flies as in the panel A, a total of 143,200 frames, frames per animal range: 2,250 –18,650). Each dot was a frame. Colormaps were applied to represent animals and stages. Blues represent clonic seizures. Reds represent tonic seizures. Oranges represent spasms. Purples represent clonic seizures. Greens represent recovery episodes. The sequential colors in each colormap represent individual animals. (**F**) Softmax classifiers were trained on flies from one line and used to decode stages in flies from other lines. Weighted F1 score was used to measure the classification accuracy.

Quantitative evaluation revealed the LASC model achieved robust stage classification, with precision, recall, and weighted F1 scores all exceeding 0.90 in the validation set after sufficient training epochs (Fig. 4B). The confusion matrix between manually-labeled and model-predicted stages showed that predicting accuracy ranging from 1.0 for paralysis to 0.97 for clonic seizures to 0.84 for spasm recognition (Fig. 4C). Based on the principle of classification, time series that have similar motion features would get the same stage label. We extracted the latent features from the attention-weighted LSTM layer (LSTM * W) and used them for visualization with uniform manifold approximation and projection (UMAP) and for quantifying seizure stage generalizability across *jus* lines (Fig. 4D). UMAP visualization revealed five distinct clusters corresponding to the seizure stages, with significant overlap between identical stages across different subjects and genotypes (Fig. 4E, Supplementary fig. 3). The shareable nature of these stages was further demonstrated through cross-genotype decoding tests, where classifiers trained on one mutant strain successfully identified stages in others with high accuracy (Fig. 4F, weighted F1 score range: 0.84-0.99). This inter-strain generalizability strongly suggests that while seizure development may vary with genetic background, the fundamental blocks of seizure progression remain consistent across *jus* alleles.

These results demonstrated: (1) a high level of discriminatory information persists in leg movement dynamics, (2) The seizure stages characterized by leg movements are shared across the four *jus* mutant lines. The LASC model enables objective prediction of seizure stages across genetic backgrounds, establishing a foundation for examining higher-order behavioral patterns.

### Quantification of seizure stages

We quantified three motion features across seizure stages: leg speed, leg movement variability, and inter-leg synchrony. Stage episodes were defined as continuous frames sharing the same classification label. Our measurements revealed distinct kinematic signatures for seizure stages, validating both our model predictions and visual observations.

Leg speed exhibited significant stage-dependent differences (Kruskal-Wallis test with Dunn’s multiple comparisons): paralysis (P) < tonic seizure (T), spasm (S), clonic seizure (C) < recovery episode (R). Recovery episodes showed the highest speeds (p = 7.6×10^-5^, 2.63×10^-10^, and 3.3×10^-5^ versus T, S, and C respectively), while paralysis demonstrated the lowest (p = 8×10^-6^, 2.91×10^-3^, and 4×10^-6^ versus T, S, and C; Fig. 5A). Movement variability, best captured by leg-body angle measurements, revealed that tonic seizures and recovery episodes showed significantly greater variation than other stages (T vs P/S/C: p = 2.23×10^-4^, 0.001003, 0.00226; R vs P/S/C: p = 5.9×10^-5^, 2.34×10^-4^, 5.46×10^-4^; Fig. 5B). This aligns with the characteristic large, stiff movements of tonic phases and the struggling motions of recovery episode, contrasting with the limited movement during paralysis, spasms and clonic seizures. Inter-leg synchrony quantification demonstrated that clonic seizures were uniquely characterized by highly synchronized leg movements (p = 4.53×10^-4^, 0.003356, 0.010054, 0.010385, C versus P/T/S/R, Fig. 5C). Notably, we found no significant bias in seizure expression across legs, as confirmed by both speed and correlation analyses (clonic seizures as a representative stage, One-way ANOVA, Fig. 5D,E).

**Figure 5.**
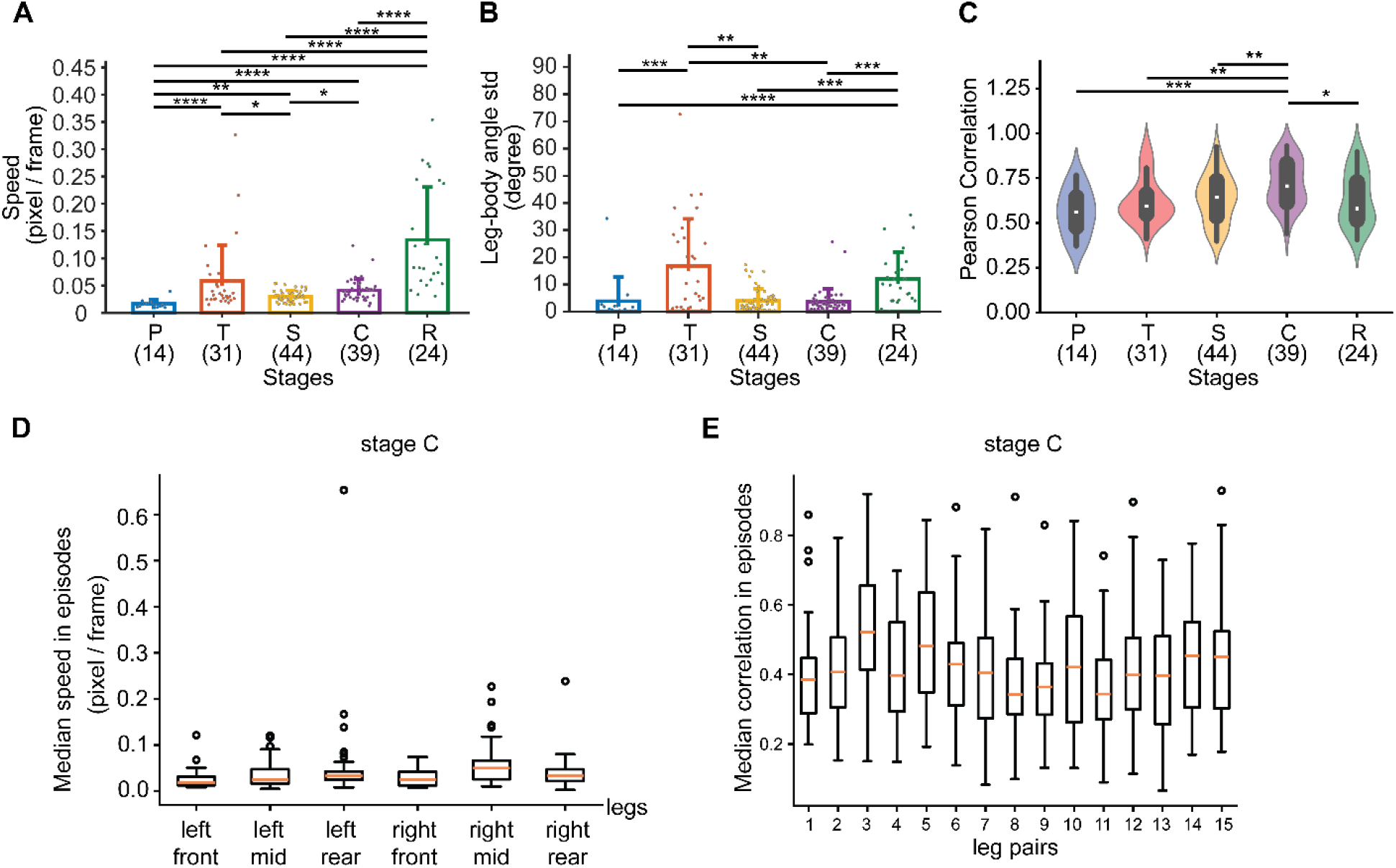
Quantitative analysis of *jus* seizure stages. (**A**) Leg speed was compared across seizure stages: P < T, S, C < R, S < T, C. Datapoints were stage episodes from the four lines. For each episode, the median value was obtained and then averaged over the six legs. Kruskal-Wallis test followed by Dunn’s multi comparison as post hoc test. *p < 0.05, **p < 0.01, ****p < 0.0001. (**B**) The variation of the leg-body angle (the angle between femur, vector from leg base to femur-tibia joint, and body-center axis, vector from abdomen tip to neck) were compared among seizure stages: P, S, C < T, R. The standard deviation per episode was obtained and then averaged over the six legs. Kruskal-Wallis test followed by Dunn’s multi comparison as post hoc test. **p < 0.01, ***p < 0.001, ****p < 0.0001. (**C**) The Pearson correlation values of leg movement were compared among seizure stages: P, T, S, R < C. For each episode, the median value was obtained and then the maximum value among the 15 leg pairs was taken. Kruskal-Wallis test followed by Dunn’s multi comparison as post hoc test. *p < 0.05, **p < 0.01, ***p < 0.001. (**D**) The seizure expression was not selectively expressed in any particular leg. Speed of legs showed no difference in the clonic seizure stage. One-way ANOVA test. (**E**) Pearson correlation of leg pairs showed no difference in the clonic seizure stage. One-way ANOVA test.

The strong concordance between these computational quantifications and visual, qualitative observations (as stated in Table 1) establishes the LASC model as a reliable framework for automated labelling of seizure stages.

### Stage sequences reveal a stereotyped progression of seizure stages

Building upon our classification of seizure stages based on leg motor patterns, we examined how these stages progress temporally across the four *jus* mutant lines. By constructing comprehensive ethograms for each genetic line, we identified a dominant seizure progression pathway that follows the sequence: paralysis (P) → tonic seizure (T) → spasm (S) → clonic seizure (C) → recovery episode (R), represented by the outermost path in the transition diagrams (Fig. 6A-D). This typical sequence emerged as the statistically most probable progression, suggesting a stereotyped architecture underlying seizure development.

**Figure 6.**
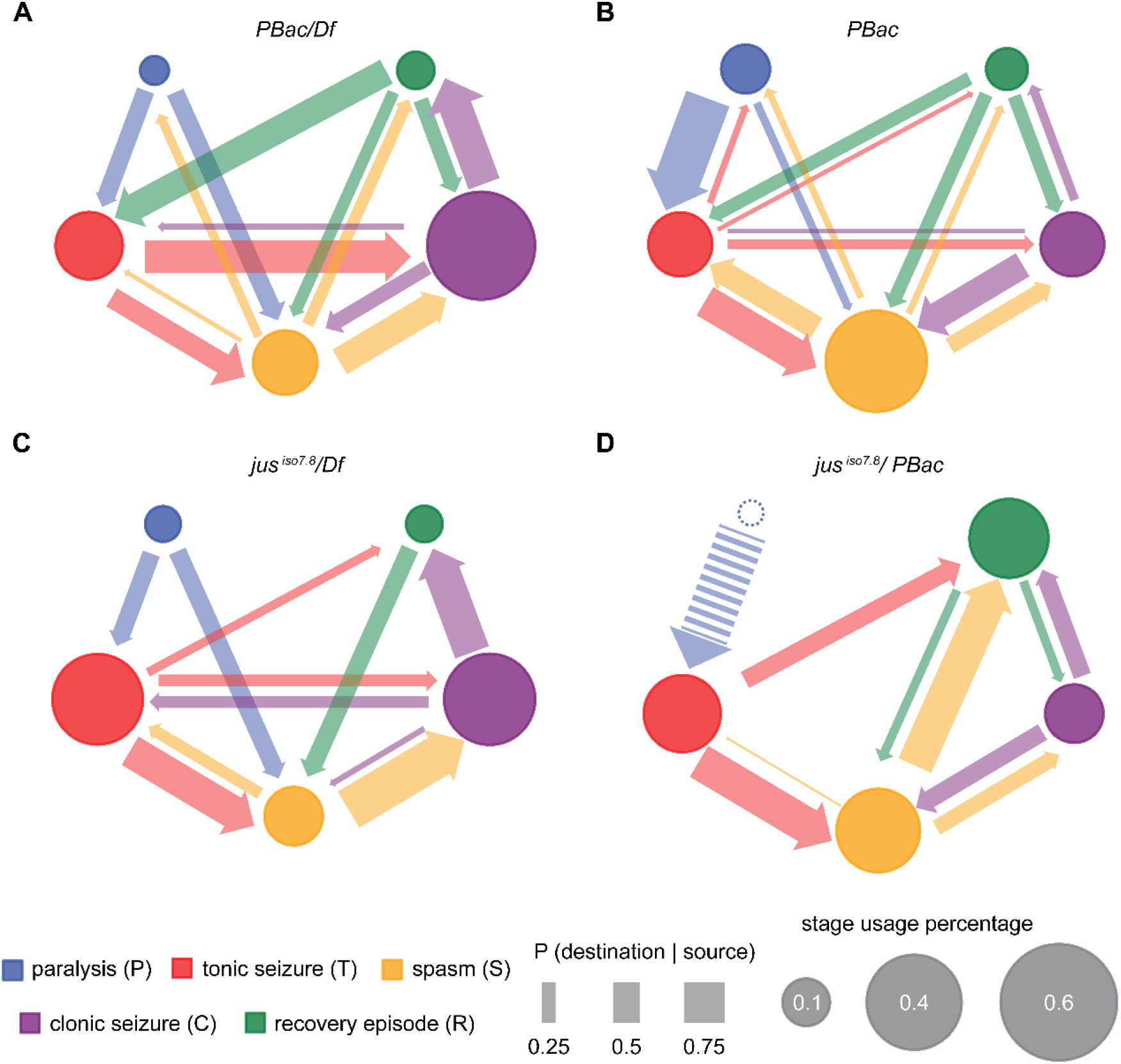
*jus* seizures show stereotyped stage transition patterns. (**A**) The ethogram illustrates different stages and the transitions performed during seizures for *PBac/Df* flies (n = 3 flies). The sizes of the nodes are proportional to the percentage of the stage usage. Arrow widths represent the transition probabilities. (**B**) The ethogram for *PBac* flies (n = 4 flies). (**C**) The ethogram for *jus* ^*iso7*.*8*^*/Df* flies (n = 3 flies). (**D**) The ethogram for *jus* ^*iso7*.*8*^*/PBac* flies (n = 3 flies). Initial stage is paralysis for each genotype (blue circle). For *jus*^*iso7*.*8*^/*PBac*, paralysis is inferred but not observed due to its brief stage.

However, seizure stage progression was not unidirectional. Closer examination revealed significant genotype-specific variations in how strictly the typical sequence was followed. The *PBac/Df* line displayed particularly frequent cycling between clonic seizures and recovery episodes, often repeating this transition multiple times before achieving full recovery (Fig. 6A). In contrast, *PBac* homozygotes showed a distinct pattern of oscillating between T and S, and S and C (Fig. 6B), while the *jus*^*iso7*.*8*^*/PBac* mutants demonstrated an increased likelihood of transitioning directly from either tonic or spasm phases into recovery episodes (Fig. 6D).

The ethograms systematically excluded full recovery states (defined by righting body and standing upright), which occurs after a recovery episode and marked the endpoint of a seizure. The ethograms collectively demonstrated that seizure progression in *jus* mutants followed a stereotyped sequence, P→ T→ S→ C→ R, but with genotype-dependent modulation of transition dynamics. The conservation of stage progression across genetic backgrounds points to deeply rooted neural circuits governing seizure development, while the observed variations in transition patterns highlight the role of genetic modifiers in regulating progression through this conserved framework.

### Seizure behavioral structures reveal distinct genotypic signatures

Next, we assessed the behavioral structures of each *jus* line. By extracting and analyzing stage sequence features – including absolute frame counts and fractional stage durations – we identified remarkable differences in how each line manifests the seizure stages (χ^2^ > 934.77, p < 4.87 × 10^-201^, Chi-squared test; Fig. 7A-B). *PBac* and *jus*^*iso7*.*8*^*/PBac* insertion lines exhibited unimodal patterns peaking at spasm (S). *PBac/Df* showed a high peak at clonic seizure (C), while *jus*^*iso7*.*8*^*/Df* displayed a bimodal distribution with peaks at both tonic (T) and clonic (C) phases. Notably, lines carrying the weak *jus*^*iso7*.*8*^ hypomorph consistently showed reduced overall seizure duration, as evidenced by smaller areas under their stage distribution curves.

**Figure 7.**
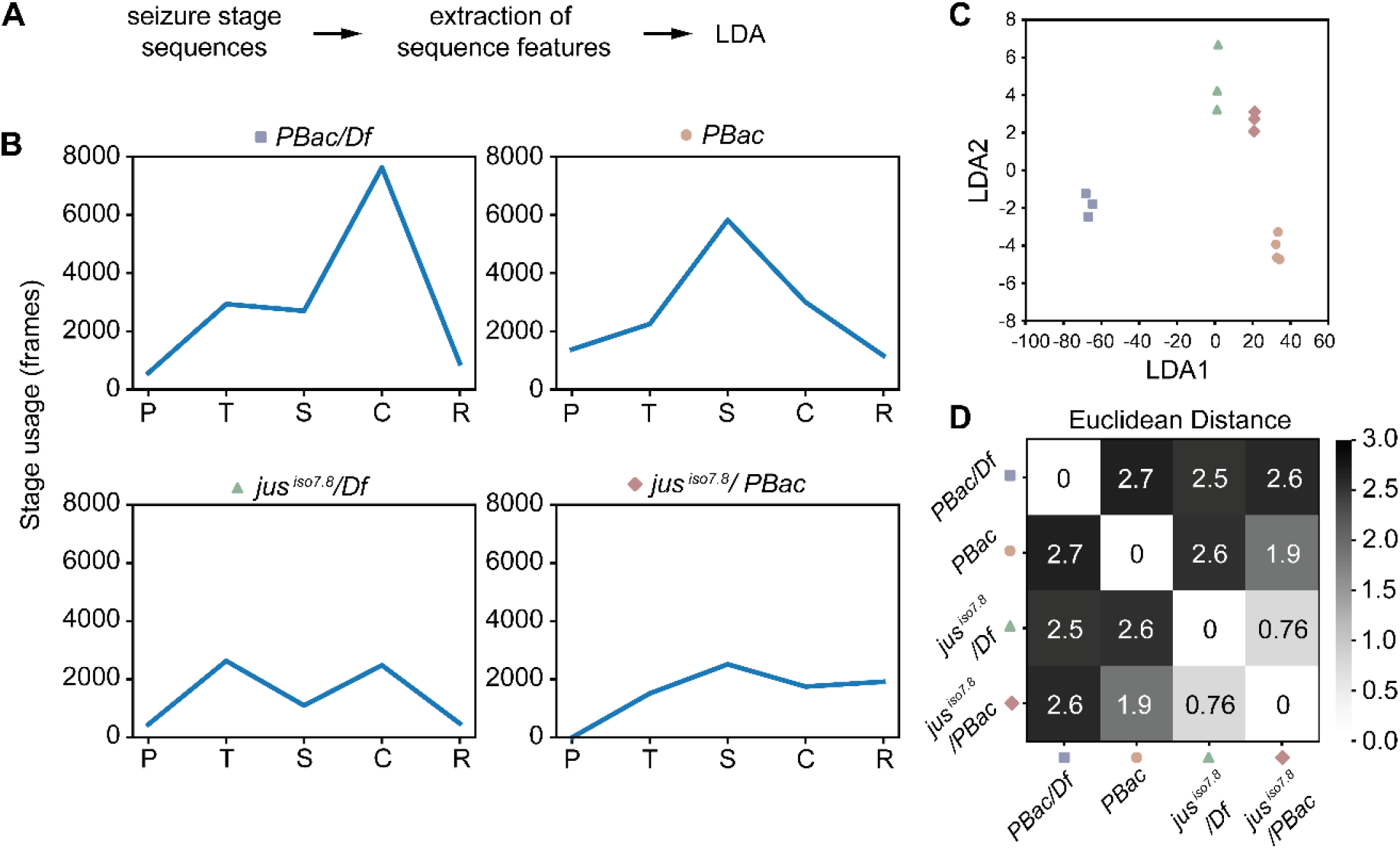
*Drosophila* genotypes are distinctive in their seizure stage usage. (**A**) Pipeline for analyzing stage sequences. (**B**) Stage usage distributions of *jus* lines. Frame numbers of each stage, ordered in paralysis (P); tonic seizure (T); spasm (S); clonic seizure (C); recovery episode (R), were shown for each line. Chi-square tests were conducted across the four lines and between every two lines to examine the difference of the usage distribution. All genotypes were significantly different from each other (χ^2^ > 934.7, p-value < 4.87e-201). (**C**) Linear discrimination analysis (LDA) indicated the differences and similarity of behavioral summaries of *Drosophila* flies. (**D**) Quantification of the behavioral differences across *jus* lines by calculating Euclidean distances based on LDA features.

To visualize the relationship between the behavioral structures and genotypes, we performed linear discriminant analysis (LDA) on the sequence features, which clustered the flies and cleanly separated all four genotypes in the resulting 2D projection (Fig. 7C). The analysis also revealed behavioral similarity: lines sharing the *jus*^*iso7*.*8*^ allele clustered closely. Euclidean distance measurements between genotype centroids confirmed that behavioral distances precisely (Fig. 7D).

These findings reveal that, although total seizure durations and individual stage lengths show minimal statistical variation across genotypes (Supplementary Fig. 4), they fail to capture the deeper distinctions encoded in the structure of seizure behavior. In contrast, the specific pattern of stage usage—defined by the relative timing and absolute duration of each phase—provides a robust, genotype-specific behavioral fingerprint. This striking correspondence between seizure dynamics and genotype indicates that *jus* mutations reshape not only seizure severity but also the underlying neural circuit activity driving seizure progression. Our results establish structured behavioral analysis as a powerful, high-resolution tool for seizure phenotyping.

## Discussion

Extensive behavioral screening in *Drosophila* has discovered mutants exhibiting the “Bang-sensitive” (BS) phenotype following mechanical stimulation ^4,8,10,18^. The seizure phenotype in *eas, tko*, and *jus*^*iso7*.*8*^ (*sda*) mutants follows a consistent four-phase progression: initial seizure (pre-paralysis hyperactivity) → paralysis → post-paralysis hyperactivity → recovery ^9,18,19^. The hyperactive phases are marked by characteristic behaviors including falling over, vigorously flapping of wings, leg shaking, and abdomen flexion. Notably, *para*^*bss1*^ mutants exhibit more severe tonic-clonic seizure activity than other BS mutants ^10,30^. While these behavioral phases have been well-documented qualitatively, current quantitative analyses remain limited to a relatively crude metric – recovery time following mechanical stimulation. Previously, we reported distinct recovery time in several *julius seizure* (*jus*) lines ^12^. However, this approach fails to capture the rich behavioral architecture underlying seizure progression. Here, our study addresses this gap by developing a comprehensive stage classification system. Through detailed stage classification and sequencing, we identified tonic seizure and clonic seizure subphases and revealed both similarities and differences of seizure progression across different *jus* lines, providing high resolution in the analysis of seizure behavioral dynamics.

Through high-speed videography and computer vision analysis, we precisely measured fine-scale movements, including the dynamics of leg joints and tips that were just 2-4 pixels in width. This enabled the identification of leg movement patterns and the corresponding five distinct behavioral stages between the mechanical stimulation and full recovery: (1) paralysis, (2) tonic seizure, (3) spasm, (4) clonic seizure, and (5) recovery episode. To systematically analyze stages, we developed an analytical pipeline centered on LASC (<u>L</u>ong short-term memory and <u>A</u>ttention mechanism for <u>S</u>equence <u>C</u>lassification), an AI model that classifies behavioral stages by analyzing spatiotemporal leg movement patterns derived from body part coordinate trajectories. Application of LASC to four distinct *jus* lines revealed several key findings: First, all mutants exhibited the same five stages, confirming conserved behavioral basis. Second, ethogram analysis demonstrated a stereotyped transition sequence (paralysis → tonic seizure → spasm → clonic seizure → recovery episode) while revealing striking genotype-specific differences in stage duration and transition patterns. Most remarkably, the genotype of an individual fly can be decoded from its seizure stage usage pattern. These findings demonstrated that: *jus* seizures follow stereotyped behavioral rules, yet manifest in genotype-specific distinctions that can serve as behavioral fingerprints. This dual observation of stereotypy and diversity in motor patterns points to both fundamental neural circuits and how genetic variation modulates neural circuit output in seizure activity.

Our behavioral analysis reveals that the progression and structure of *jus*-mediated seizures are fundamentally shaped by genetic background. This genotype dependence raises two pivotal questions about the neural basis of seizure behavior. First, how does the CNS essentially encode the distinct seizure stages? Second, what is the mechanistic link between *jus* expression and the resulting neural dynamics? It revealed an intriguing pattern when we examine seizure episodes more closely: each seizure episode may involve different legs (Fig. 2), but when we examined all episodes with the same stage label, they showed no leg preference (Fig. 5). This variability leads us to a third question: how do motor circuits differentially engage legs during seizures? From the known neural circuit, leg movements are precisely controlled by motor neurons in the thoracic-abdominal ganglion (TAG) ^31^. Intriguingly, *jus* is expressed throughout this system—most prominently in the first two TAG segments, but also in brain regions like the optic and antennal lobes ^11^. Previous work established that *jus* plays particularly crucial roles in cholinergic and GABAergic neurons, where it acts as a seizure threshold gatekeeper ^11,12^. These findings indicated the complex relationship between genes, neural circuits and behaviors. We propose that spatial and temporal patterns of *jus* expression regulate neuronal excitability and seizure thresholds, thereby governing the emergent dynamics of seizure networks. The *jus* seizure model offers a powerful experimental platform for systematically dissecting the causal chain of gene expression patterns, neural circuit dynamics, and stage-specific behavioral outputs. Mapping behavior to neural activities has been a fundamental objective in neuroscience and serves as a crucial step toward understanding how the brain drives behavior ^32,33^. Indeed, mapping these relationships presents challenges. While modern imaging tools allow us to monitor specific neuronal populations in the brain ^34^ or the ventral nerve cord ^35,36^ with impressive precision, the fly’s small size makes simultaneous behavior-neural recording notoriously difficult. Reliable imaging approaches require physical fixation, which is ideal for imaging stability but incompatible with vortex-induced seizures. Here, *jus* mutants provide a solution through cold shock induction. Consistent seizure phenotypes can be reliably triggered by brief (1-minute) low-temperature exposure^11^. This alternative induction modality offers two key advantages: compatibility with fixed preparations for stable neural recordings, and activation through distinct sensory pathways compared to mechanical induction. Notably, advances in imaging and computational analysis of neural dynamics could be powerfully combined with this model to dissect the gene regulated neural mechanisms behind seizure behavioral phenotypes.

Beginning with raw video data, we progressively extract and organize movement information across multiple scales – from individual body part kinematics to integrated action sequences to behavior spectrum – creating a comprehensive behavioral embedding space. This hierarchical approach captures both the subtle movement signatures that distinguish specific stages and the broad architecture of seizure, achieving much higher resolution compared to traditional recovery-time metrics ^9,17,37–39^. The power of this framework becomes particularly evident when applied to drug screening in our *jus* mutant model ^39,40^. Where conventional bang-sensitive assays could only measure crude endpoints like recovery time, our system can detect nuanced drug effects on specific seizure phases and transition patterns. Notably, the Jus protein interacts in a complex of conserved proteins found to affect seizures, including the alpha and beta subunits of the Na^+^/K^+^ ATPase ^41^. It is possible that therapeutics modulating the homologous mammalian complex can be designed and tested to alleviate seizures in *jus* mutants ^42^. What makes this behavior-based approach particularly valuable for translational research is its combination of *Drosophila’s* practical advantages – rapid life cycle, genetic tractability, and reliable seizure induction – with analytical depth. The system enables high-throughput compound screening while maintaining the resolution needed to identify subtle modulators of seizure dynamics. As we continue to refine this platform, it promises to accelerate both the discovery of novel therapeutics and our fundamental understanding of how a gene can modulate neural circuits and control seizure

behaviors.

## Methods

### Fly stocks

Fly stocks were kept on agar media containing glucose, yeast, and cornmeal at 22 to 25°C in plastic vials. Fly stocks were obtained from the Bloomington Stock Center at Indiana University. The *jus* ^*iso7*.*8*^ mutant was a gift from M. A. Tanouye. *jus* alleles were acquired by crossing and screening. For *PBac/Df* flies, virgin females from *PBac[WH]jus*^*f04904*^ (BL#18817, a *jus* insertion line) were crossed to males from *Df(3R)BSC500* (BL#25004, a *jus* deficiency line). For *jus* ^*iso7*.*8*^*/Df* flies, virgin females from line *jus* ^*iso7*.*8*^ were crossed to males from *Df(3R)BSC500*. For *jus* ^*iso7*.*8*^*/PBac* flies, virgin females from line *jus* ^*iso7*.*8*^ were crossed to males from *PBac[WH]jus*^*f04904*^.

### Fly preparation for video recording

Visual inspection revealed that *jus* seizures manifested primarily in the legs rather than the wings. To enhance leg-background contrast for video analysis, we removed the wings prior to seizure induction and recording. Within 24 hours of adult emergence, flies were anesthetized with CO2 and screened by genotype. Selected flies underwent wing excision (Fig. 1A). Processed flies were transferred to new vials containing food and allowed to recover for ≥24 hours before experimentation.

### Seizure induction and behavior recording

On recording day, individual flies were transferred to empty vials using a pipette system. Flies were allowed to recover for at least 0.5 hour after transfer (Fig. 1B). Then, seizures were induced by vortex mixing the vial for 10 seconds using the Vortex Genie 2 (VWR Scientific, Radnor, PA) at maximum speed (Fig. 1C). Immediately post-induction, the knocked down fly was positioned in the recording chamber, affixed dorsum-down using laboratory tape, and the chamber was immediately placed in the imaging arena (Fig. 1D). This post-induction transfer typically took about 40 seconds. The video recording began right after the chamber was placed and did not stop until the fly recovered from seizure, which was indicated by the fly turning over and standing up.

### Behavior recording apparatus

The behavior recording setup consisted of an imaging chamber, LED lights and a high-speed camera (Fig. 1E). The imaging chamber was constructed with a white frosted glass base to minimize light reflection and optimize contrast. It also incorporated double-sided tape (type 415, 3M, USA) for affixing the fly at a relatively consistent recording angle. DC-powered LED array (IvisiiG2) provided stable, flicker-free illumination, which is critical for high-speed imaging. A FLIR Blackfly S BFS-U3-13Y3M USB camera equipped with a Computar 25 mm f1.4 lens and an EX2C extender for C-mount was used to record video files at 100 frames per second.

### DeepLabCut for body part tracking

We developed a video analysis pipeline (Fig. 3A). The first step is using DeepLabCut (DLC) framework to track 31 anatomically defined body parts (Supplementary Fig. 1). The analysis focused exclusively on female flies because male genitalia reduced contrast between legs and abdomen, compromising tracking accuracy. As it is a challenge to track multiple legs which are small and move in an extremely flexible manner, we chose resnet-152 as the network type. By visual inspection, most of the time, legs were accurately tracked. However, in the tonic stage, sudden and violent movements and leg crossings caused leg misidentification. We performed rounds of refinements by manually correcting the outlier frames and adding to the training set. Ultimately, the DLC model was trained with a total of 1,232 high-quality manually annotated frames. The model achieved high performance on the training set (3.8 mean pixel error for all 31 body parts and 5.17 mean pixel error for tibia-tarsus joints and leg tips) and testing set (4.07 mean pixel error for all 31 body parts and 5.53 for tibia-tarsus joints and leg tips). To minimize jitter in the positions, we applied a median filter (window length = 5 frames) before further analysis.

### Extraction of motion features

While body part tracking greatly reduced data dimensionality from raw videos and captured behavioral dynamics, additional motion features were necessary to fully characterize seizure-related actions. We focused on leg movement dynamics, deriving 159 quantitative features from tracked XY coordinates (Supplementary Table 1 for complete list and formulas). Key feature categories included: (1) corrected distance of each leg tip; (2) speed of each leg tip; (3) acceleration of each leg tip; (4) angle of the tibia-tarsus joint; (5) angular speed of the tibia-tarsus joint, (6) the angle between femur (vector from leg base to femur-tibia joint) and body-center axis (vector from abdomen tip to neck); (7) Pearson correlation R between every two leg tips by using features of corrected distance, X and Y coordinates, speed and acceleration. R were calculated in a sliding window and centered (window size = 50 frames). These motion features captured kinetic, postural and inter-leg synchronization properties.

### LASC model for seizure stage classification

We developed a <u>L</u>ong short-term memory and <u>A</u>ttention mechanism for <u>S</u>equence <u>C</u>lassification (LASC) method to classify five distinct *jus* seizure stages: paralysis, tonic seizure, spasm, clonic seizure, and recovery episode. 159 motion features with a time step of 50 frames were used as input and each input was labelled with one of the predefined five stages. The labelled dataset was randomly split for training (70%), validation (15%) and test (15%). In total, 2202 inputs (110,100 frames) were used in the model training. LASC is based on long-short term memory (LSTM) algorithm. The layer architecture was shown in Fig. 3D. The number of units of the LSTM layer was fixed to 64. The LSTM layer was followed by an attention layer to apply weights on the LSTM latent features. Then the outputs were summed over the time dimension. In the end, a softmax layer was used for classification. The network was trained using weighted categorical cross-entropy loss and the Adam optimizer in TensorFlow 2.10.0. The model achieved good performance on the test set (precision = 0.9083, recall = 0.9025). We adopted the trained LASC model to implement behavioral sequencing. In the four lines (Fig. 4A), 92850 frames (64.8% of the total frames in these videos) were used for the LASC model training or evaluation. To reduce noise, the predicted stage labels were smoothed using a sliding-window mode over 250 frames.

### Model comparison and performance assessment

We conducted a comprehensive performance evaluation comparing our LASC model against three deep learning architectures commonly used for time-series classification: a basic LSTM without attention mechanisms, a CNN, and a transformer model. The layer structures for each model can be found in the Supplementary Table 2. All models shared the same training dataset, ensuring a fair comparison of their ability to classify five distinct behavioral stages. For training, we applied the Adam optimizer with the dropout or L_2_ norm regularization methods. We use weighted categorical cross-entropy as the loss function, which scaled the loss by class weights to account for class imbalances. The training process incorporated early stopping (patience=15 epochs, maximum 100 epochs) to optimize model convergence. We explored sets of parameters for each model and used the one that performed the best on the validation dataset to implement unbiased comparison on the reserved test dataset. For performance evaluation, we used precision, recall and weighted F1 score. For each stage, precision was calculated by: TP / (TP + FP), where TP means true positives (samples are correctly predicted as the stage label), and FP means false positives (samples are incorrectly predicted the stage label). Recalls are calculated by TP / (TP + FN), where FN means false negatives (samples of the stage are incorrectly predicted as other classes). The weighted F1-score, calculated as the harmonic mean of precision and recall 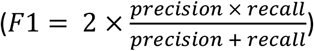 averaged across classes proportionally to their sample sizes, provided our primary measure of classification accuracy. Our results demonstrated that the LASC model outperformed the others with consistently higher precision, recall, and weighted F1-scores. This performance advantage on each stage is visually apparent in the confusion matrices presented in Fig. 4C and Supplementary Fig. 2B-D. The attention mechanism in LASC proved particularly valuable for behavioral classification, enabling the model to dynamically focus on the most informative features in each sequence while maintaining awareness of long-range temporal dependencies. This architecture allowed LASC to better capture the nuanced differences between behavioral stages compared to the standard LSTM, CNN, and transformer models, ultimately yielding more accurate and reliable classification of complex behaviors.

### Uniform Manifold Approximation and Projection (UMAP) embedding

We used the UMAP technique to embed the behavioral latent features in low dimensional space. Latent features were extracted from our LASC model (attention-weighted layer in Fig. 3D, LSTM * W). UMAP visualization can provide intuition of the behavioral repositories. We performed UMAP analysis on datasets of combined animals. The same UMAP reducer was adopted for all analysis. The parameters that we used were (n_components = 2, n_neighbors = 200, min_dist = 0.5, random_state = 52, others = default). We explored varying the n_neighbors and min_dist parameters and obtained the optimized results.

### Cross-line decoding analysis

To assess seizure stage generalizability across *jus* lines, we performed cross-line decoding experiments. We trained a softmax classifier on latent features from a source *jus* line and evaluated its performance on other target *jus* lines. As for the source *jus* line, we randomly split its labeled dataset into training (85%) and validation (15%) sets. As for the target *jus* lines, we used the entire labeled dataset for testing.

### Quantification of stages

Using the predicted stage labels, we segmented the time-series data into episodes of continuous identical-stage frames for motion feature quantification. For each feature type, we employed the following analytical approaches: 1. Speed comparison: Calculated the median speed per episode, then averaged across all six legs. 2. Leg-body angle variability: Determined the standard deviation per episode, then averaged across all six legs. 3. Inter-leg synchronization: Used Pearson correlation coefficients of corrected distance, taking the maximum value among all 15 leg pairs (6 choose 2) for each episode.

### Ethogram construction

Based on the predicted stage labels, we analyzed the behavioral transitions. First, we computed raw transition counts between stages. Then, the counts of each transition were divided by the total occurrence of the corresponding starting stage to obtain the transition probabilities. The ethogram (Fig. 6) was constructed to describe the frequency of each behavioral stage and the probability that one stage is followed by another. The sizes of the nodes were proportional to stage occurrence frequency. Line widths represented the transition probabilities between the stages.

### Linear discrimination analysis (LDA) and Euclidean distance

To assess similarities and differences across genotypes, we embedded behavioral summaries into low-dimensional space. First, stage usage features were extracted from behavioral sequences. Both absolute frame count (*stage_usage_frameNum*) and relative percentage (*stage_usage_pct*) were used. Then, linear discrimination analysis (LDA, scikit-learn package in Python) was used to calculate a projection of the stage usage features, which maximized linear separability between groups. The input of LDA analysis in Fig. 7C was (13 seizure sessions × standardized 10 stage usage features). To quantify the distance of groups in the LDA space, pairwise Euclidean distance was computed (sklearn package in Python). Two LDA features were standardized and then averaged within groups. The Euclidean distance between groups was calculated and shown in a symmetric matrix.

### Statistics

Statistical analyses were performed in Python with appropriate methods as indicated in results and figure legends. Unless otherwise specified, data were shown as mean ± sd. We did not use statistical methods to predetermine sample size. No outliers or other data were excluded. For all analysis, a p-value < 0.05 was considered statistically significant. The significance is indicated as * p < 0.05, ** p < 0.01, *** p < 0.001 and **** p < 0.0001. The details of statistical tests and results were shown in the results and figure legends. All figures were prepared using Python and Illustrator CC (Adobe).

## Data availability

All raw video recordings in this study are available from the corresponding author upon reasonable request.

## Code availability

All custom codes that support the findings of this study are available at:

https://github.com/glaulab/LASC, which includes the motion feature extraction, LASC model construction and compilation, visualization of the behavioral sequence data, and video processing.

## Acknowledgements

This work was supported by the Hong Kong Research Grants Council (RGC/GRF 11104521 to C.G.L.), internal funds from City University of Hong Kong (9610354 and 9380104 to C.G.L.), and the Jockey Club College of Veterinary Medicine and Life Sciences of City University of Hong Kong / Cornell University Graduate Program Fund (D.L.D.). We thank Nilay Yapici (Cornell) for help with setting up the high-speed camera and Itai Cohen (Cornell) for help with improving the behaviour recordings platform. We thank Runpeng HOU, Huilin ZHAO and Arthur CHIU (CityUHK) for discussions during the early stages of this work.

## Author contributions

H.Z. performed the experiments, analyzed data and wrote the first draft of the manuscript. C.G.L. and D.L.D reviewed and revised the manuscript. C.G.L., D.L.D and H.Z. conceived the study. C.G.L. and D.L.D supervised the study and obtained funding. All authors read and approved the final manuscript.

## Supplementary information

Supplementary tables and figures are included in a file.

