## Supplemental tables and figures for "Automated behavior classification of *julius seizure* mutants in *Drosophila* reveals stereotyped seizure stages with genotype specificity"

### Supplementary Materials

#### Supplementary Table 1. Motion feature list. - (Please find it at the end of the document)

#### Supplementary Table 2. Layer structures for LASC, standard LSTM, CNN and transformer classifiers.

| | Layer | Output Shape | Parameters/Notes |
|---|---|---|---|
| **LASC (LSTM + attention)** | Input Layer | (None, 50, 159) | Raw time-series input (50 timesteps × 159 features) |
| | LSTM Layer | (None, 50, 64) | units=64, return_sequences=True, dropout=0.2, recurrent_dropout=0.2 |
| | Transpose | (None, 64, 50) | Swaps axes for feature attention |
| | Feature Attention | (None, 64, 50) | Dense(1, activation='tanh'), Softmax, Multiply |
| | Transpose Back | (None, 50, 64) | Restores original shape |
| | Sum Pooling | (None, 64) | Aggregates across timesteps |
| | Output Layer | (None, 5) | softmax activation for classes |
| **LSTM (no attention)** | Input Layer | (None, 50, 159) | Raw time-series input (50 timesteps × 159 features) |
| | LSTM Layer | (None, 50, 64) | units=64, return_sequences=True, dropout=0.2, recurrent_dropout=0.2 |
| | Sum Pooling | (None, 64) | Aggregates across timesteps |
| | Output Layer | (None, 5) | softmax activation for classes |
| **CNN** | Input Layer | (None, 50, 159) | Raw time-series input (50 timesteps × 159 features). |
| | Conv1D (1st) | (None, 48, 64) | filters=64, kernel_size=3, activation='relu' |
| | MaxPooling1D (1st) | (None, 24, 64) | pool_size=2 |
| | Conv1D (2nd) | (None, 22, 32) | filters=32, kernel_size=3, activation='relu' |
| | MaxPooling1D (2nd) | (None, 11, 32) | pool_size=2 |
| | Flatten | (None, 352) | Converts the output to 1D for the dense layer. |
| | Dense (Hidden) | (None, 100) | units=100, activation='relu' |
| | Dropout | (None, 100) | rate=0.5 |
| | Dense (Output) | (None, 5) | softmax activation for classes |
| **Transformer** | Input | (None, 50, 159) | Raw time-series input (50 timesteps × 159 features) |
| | PositionalEmbedding | (None, 50, 159) | Adds timestep positions |
| | MultiHeadAttention | (None, 50, 159) | 4 heads, key_dim=64 |
| | Dropout | (None, 50, 159) | Rate=0.2 |
| | LayerNormalization | (None, 50, 159) | epsilon=1e-6 |
| | Dense (FFN) | (None, 50, 128) | units=128, activation=ReLU |
| | Dropout | (None, 50, 128) | Rate=0.2 |
| | LayerNormalization | (None, 50, 128) | epsilon=1e-6 |
| | GlobalAveragePooling1D | (None, 128) | Reduces timesteps via averaging |
| | Dense (Output) | (None, 5) | softmax activation for classes |

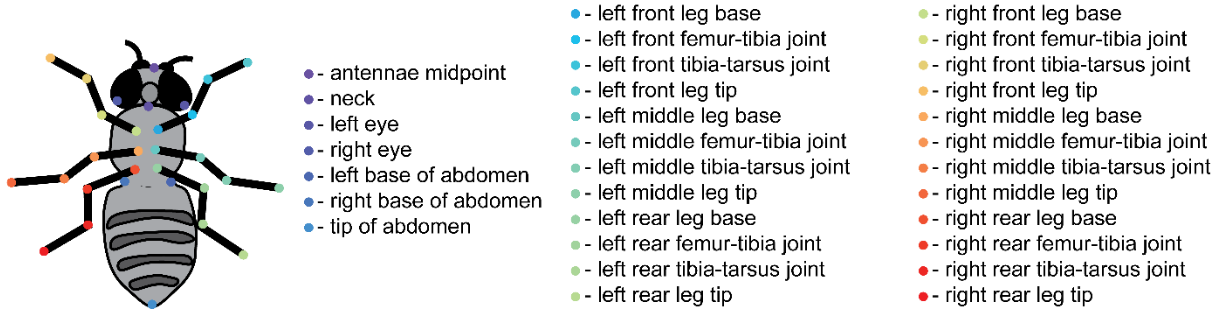

**Supplementary Figure 1. 31 Body parts in DeepLabCut (DLC) tracking.**

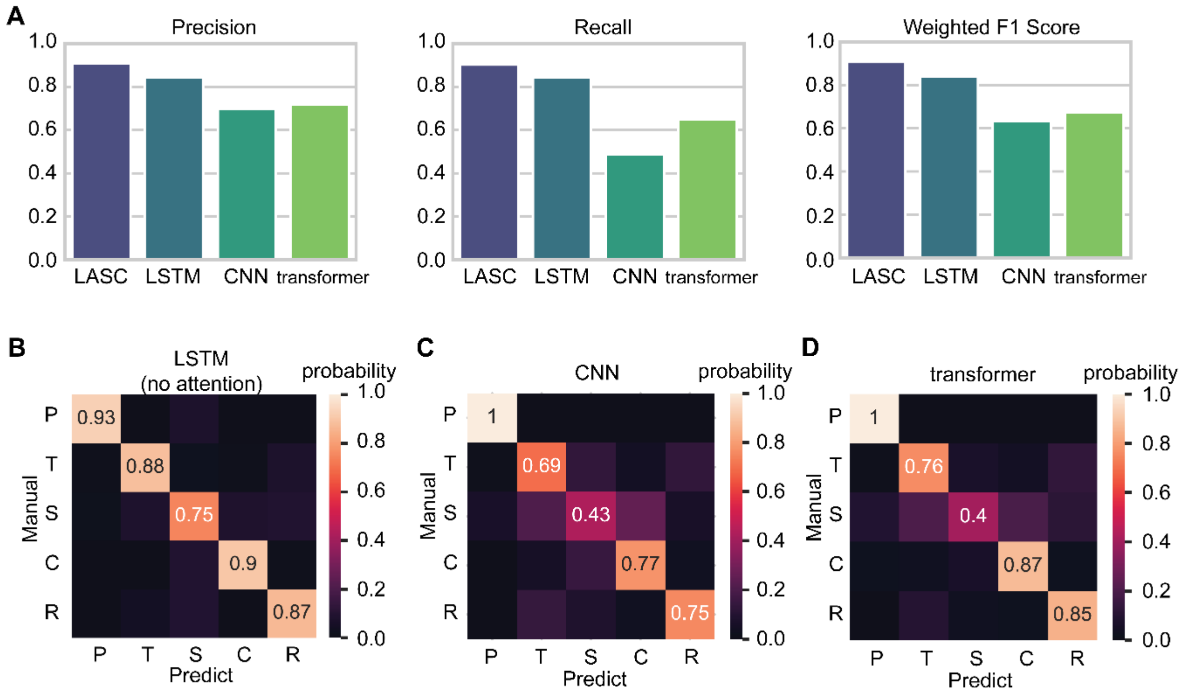

**Supplementary Figure 2. Compare LASC model with other machine learning models.** (**A**) compare models by precision, recall and weighted F1 scores. To guarantee an unbiased comparison, all models were trained on the same training dataset and tested on the same held-out test set. (**B-D**) Confusion matrix between manually labelled ground truth and model predictions. The values indicate what fraction of manually labelled data were classified as each stage by the model in the test dataset. (B) for LSTM (without attention). (C) for CNN. (D) for Transformer.

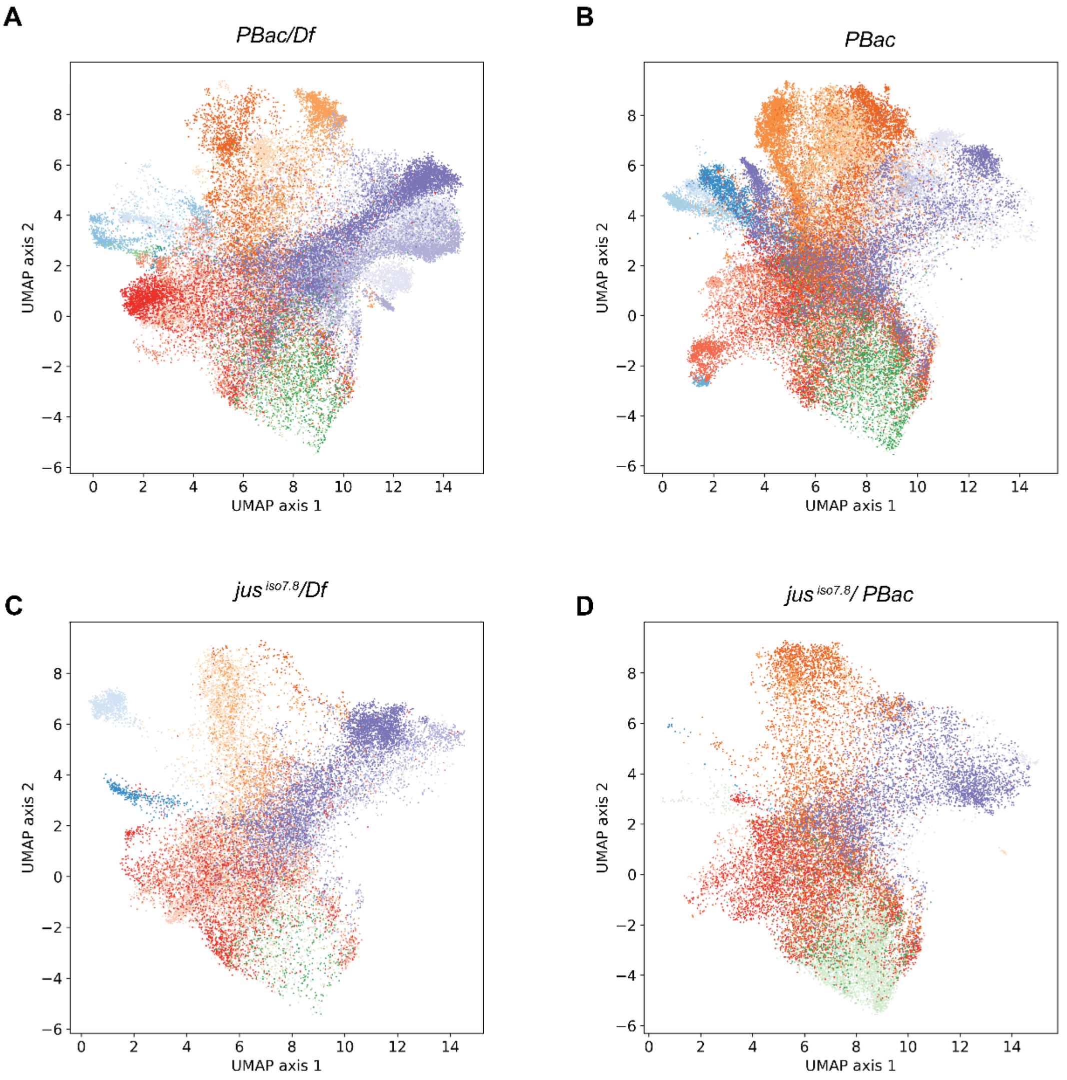

**Supplementary Figure 3. UMAP embeddings of each *jus* allele.** (**A**) UMAP embedding of LASC latent features from *PBac/Df* flies (a total of 44,200 frames, frames per animal range: 13,500 – 15,600). Color coding was the same as in Figure 4. The sequential colors represent individual animals. (**B**) UMAP embedding of LASC latent features from *PBac* flies (a total of 54,450 frames, frames per animal range: 9,950 – 18,650). (**C**) UMAP embedding of LASC latent features from *jus* $^{iso7.8}$*/Df* flies (a total of 21,450 frames, frames per animal range: 6,050 – 8,450). (**D**) UMAP embedding of LASC latent features from *jus* $^{iso7.8}$*/PBac* flies (a total of 23,100 frames, frames per animal range: 2,250 – 13,200).

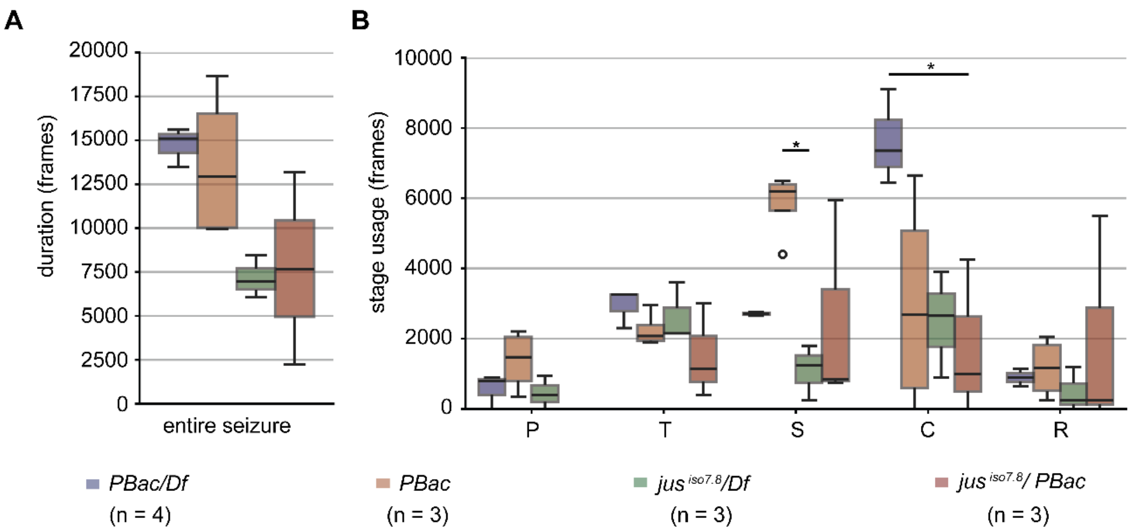

**Supplementary Figure 4. Comparisons of seizure duration and stage usage.** (**A**) Comparison of entire seizure duration among four *jus* lines. One-way ANOVA. (**B**) Comparison of stage usage among four *jus* alleles. One-way ANOVA with Tukey multiple comparison as post hoc test. *$p < 0.05$.

| Supplementary Table 1. Motion feature list | | |
|---|---|---|
| **Fearure NO.** | **Feature name** | **DLC tracking data were saved as (x, y) label position in pixel for each body part per frame.<br>n represents the total frame number of one video.<br>All features are time series. The calculation at frame, f, was shown.** |
| 1 | dist_left_front_leg_tip | corrected distance<br>$dist(f) = \sqrt{x_f^2 + y_f^2} - \frac{1}{n}\sum_{f=1}^{n}\sqrt{x_f^2 + y_f^2}$ |
| 2 | dist_left_middle_leg_tip | |
| 3 | dist_left_rear_leg_tip | |
| 4 | dist_right_front_leg_tip | |
| 5 | dist_right_middle_leg_tip | |
| 6 | dist_right_rear_leg_tip | |
| 7 | posX_left_front_leg_tip | Original tracking x and y |
| 8 | posX_left_middle_leg_tip | |
| 9 | posX_left_rear_leg_tip | |
| 10 | posX_right_front_leg_tip | |
| 11 | posX_right_middle_leg_tip | |
| 12 | posX_right_rear_leg_tip | |
| 13 | posY_left_front_leg_tip | |
| 14 | posY_left_middle_leg_tip | |
| 15 | posY_left_rear_leg_tip | |
| 16 | posY_right_front_leg_tip | |
| 17 | posY_right_middle_leg_tip | |
| 18 | posY_right_rear_leg_tip | |
| 19 | speed_left_front_leg_tip | speed<br>$speed(f) = \sqrt{(x_f - x_{f-1})^2 + (y_f - y_{f-1})^2}$ |
| 20 | speed_left_middle_leg_tip | |
| 21 | speed_left_rear_leg_tip | |
| 22 | speed_right_front_leg_tip | |
| 23 | speed_right_middle_leg_tip | |
| 24 | speed_right_rear_leg_tip | |
| 25 | acc_left_front_leg_tip | acceleration<br>$acc(f) = speed(f) - speed(f-1)$ |
| 26 | acc_left_middle_leg_tip | |
| 27 | acc_left_rear_leg_tip | |
| 28 | acc_right_front_leg_tip | |
| 29 | acc_right_middle_leg_tip | |
| 30 | acc_right_rear_leg_tip | |
| 31 | angleTTJ_left_front_leg | angle of the tibia-tarsus joint<br>$angleTTJ = \arccos(\frac{a \cdot b}{\lvert a\rvert\ \lvert b\rvert})$<br>The equation is applied for each frame,f. *a* is the vector from the tibia-tarsus joint (TTJ) to the femur-tibia joint (FTJ) and *b* is the vector from TTJ to leg tip. |
| 32 | angleTTJ_left_middle_leg | |
| 33 | angleTTJ_left_rear_leg | |
| 34 | angleTTJ_right_front_leg | |
| 35 | angleTTJ_right_middle_leg | |
| 36 | angleTTJ_right_rear_leg | |
| 37 | angSpeedTTJ_left_front_leg | angular speed of the tibia-tarsus joint<br>$angSpeedTTJ(f) = angleTTJ(f) - angleTTJ(f-1)$ |
| 38 | angSpeedTTJ_left_middle_leg | |
| 39 | angSpeedTTJ_left_rear_leg | |
| 40 | angSpeedTTJ_right_front_leg | |
| 41 | angSpeedTTJ_right_middle_leg | |
| 42 | angSpeedTTJ_right_rear_leg | |
| 43 | legBodyAngle_left_front_leg | the angle between femur and body-center axis<br>$legBodyAngle = \arccos(\frac{a \cdot b}{\lvert a\rvert\ \lvert b\rvert})$<br>The equation is applied for each frame,f. *a* is the vector from the leg base to the femur-tibia joint. *b* is the vector from abdomen tip to neck. |
| 44 | legBodyAngle_left_middle_leg | |
| 45 | legBodyAngle_left_rear_leg | |
| 46 | legBodyAngle_right_front_leg | |
| 47 | legBodyAngle_right_middle_leg | |
| 48 | legBodyAngle_right_rear_leg | |
| 49 | r_dist_left_front_leg_tip_left_middle_leg_tip | |
| 50 | r_dist_left_front_leg_tip_left_rear_leg_tip | |
| 51 | r_dist_left_front_leg_tip_right_front_leg_tip | |

| | | |
|---|---|---|
| 52 | r_dist_left_front_leg_tip_right_middle_leg_tip | |
| 53 | r_dist_left_front_leg_tip_right_rear_leg_tip | |
| 54 | r_dist_left_middle_leg_tip_left_rear_leg_tip | |
| 55 | r_dist_left_middle_leg_tip_right_front_leg_tip | |
| 56 | r_dist_left_middle_leg_tip_right_middle_leg_tip | |
| 57 | r_dist_left_middle_leg_tip_right_rear_leg_tip | |
| 58 | r_dist_left_rear_leg_tip_right_front_leg_tip | |
| 59 | r_dist_left_rear_leg_tip_right_middle_leg_tip | |
| 60 | r_dist_left_rear_leg_tip_right_rear_leg_tip | |
| 61 | r_dist_right_front_leg_tip_right_middle_leg_tip | |
| 62 | r_dist_right_front_leg_tip_right_rear_leg_tip | |
| 63 | r_dist_right_middle_leg_tip_right_rear_leg_tip | |
| 64 | r_posX_left_front_leg_tip_posX_left_middle_leg_tip | |
| 65 | r_posX_left_front_leg_tip_posX_left_rear_leg_tip | |
| 66 | r_posX_left_front_leg_tip_posX_right_front_leg_tip | |
| 67 | r_posX_left_front_leg_tip_posX_right_middle_leg_tip | |
| 68 | r_posX_left_front_leg_tip_posX_right_rear_leg_tip | |
| 69 | r_posX_left_front_leg_tip_posY_left_front_leg_tip | |
| 70 | r_posX_left_front_leg_tip_posY_left_middle_leg_tip | |
| 71 | r_posX_left_front_leg_tip_posY_left_rear_leg_tip | |
| 72 | r_posX_left_front_leg_tip_posY_right_front_leg_tip | |
| 73 | r_posX_left_front_leg_tip_posY_right_middle_leg_tip | |
| 74 | r_posX_left_front_leg_tip_posY_right_rear_leg_tip | |
| 75 | r_posX_left_middle_leg_tip_posX_left_rear_leg_tip | |
| 76 | r_posX_left_middle_leg_tip_posX_right_front_leg_tip | |
| 77 | r_posX_left_middle_leg_tip_posX_right_middle_leg_tip | |
| 78 | r_posX_left_middle_leg_tip_posX_right_rear_leg_tip | |
| 79 | r_posX_left_middle_leg_tip_posY_left_front_leg_tip | |
| 80 | r_posX_left_middle_leg_tip_posY_left_middle_leg_tip | |
| 81 | r_posX_left_middle_leg_tip_posY_left_rear_leg_tip | |
| 82 | r_posX_left_middle_leg_tip_posY_right_front_leg_tip | |
| 83 | r_posX_left_middle_leg_tip_posY_right_middle_leg_tip | |
| 84 | r_posX_left_middle_leg_tip_posY_right_rear_leg_tip | |
| 85 | r_posX_left_rear_leg_tip_posX_right_front_leg_tip | |
| 86 | r_posX_left_rear_leg_tip_posX_right_middle_leg_tip | |
| 87 | r_posX_left_rear_leg_tip_posX_right_rear_leg_tip | |
| 88 | r_posX_left_rear_leg_tip_posY_left_front_leg_tip | |
| 89 | r_posX_left_rear_leg_tip_posY_left_middle_leg_tip | |
| 90 | r_posX_left_rear_leg_tip_posY_left_rear_leg_tip | |
| 91 | r_posX_left_rear_leg_tip_posY_right_front_leg_tip | |
| 92 | r_posX_left_rear_leg_tip_posY_right_middle_leg_tip | |
| 93 | r_posX_left_rear_leg_tip_posY_right_rear_leg_tip | |
| 94 | r_posX_right_front_leg_tip_posX_right_middle_leg_tip | |
| 95 | r_posX_right_front_leg_tip_posX_right_rear_leg_tip | |
| 96 | r_posX_right_front_leg_tip_posY_left_front_leg_tip | |
| 97 | r_posX_right_front_leg_tip_posY_left_middle_leg_tip | |
| 98 | r_posX_right_front_leg_tip_posY_left_rear_leg_tip | |
| 99 | r_posX_right_front_leg_tip_posY_right_front_leg_tip | |
| 100 | r_posX_right_front_leg_tip_posY_right_middle_leg_tip | |
| 101 | r_posX_right_front_leg_tip_posY_right_rear_leg_tip | The absolute value of Pearson correlation of two time series (features) in a rolling window were calculated by using pandas package in Python.<br><br>rolling_r = df1.rolling(window=50, center=True).corr(df2)<br>rolling_r = rolling_r.abs()<br>Shown as the feature name, df1 is one time series of a feature, and df2 is another time series of a feature. |
| 102 | r_posX_right_middle_leg_tip_posX_right_rear_leg_tip | |
| 103 | r_posX_right_middle_leg_tip_posY_left_front_leg_tip | |
| 104 | r_posX_right_middle_leg_tip_posY_left_middle_leg_tip | |
| 105 | r_posX_right_middle_leg_tip_posY_left_rear_leg_tip | |
| 106 | r_posX_right_middle_leg_tip_posY_right_front_leg_tip | |
| 107 | r_posX_right_middle_leg_tip_posY_right_middle_leg_tip | |
| 108 | r_posX_right_middle_leg_tip_posY_right_rear_leg_tip | |
| 109 | r_posX_right_rear_leg_tip_posY_left_front_leg_tip | |

| | |
|---|---|
| 110 | r_posX_right_rear_leg_tip_posY_left_middle_leg_tip |
| 111 | r_posX_right_rear_leg_tip_posY_left_rear_leg_tip |
| 112 | r_posX_right_rear_leg_tip_posY_right_front_leg_tip |
| 113 | r_posX_right_rear_leg_tip_posY_right_middle_leg_tip |
| 114 | r_posX_right_rear_leg_tip_posY_right_rear_leg_tip |
| 115 | r_posY_left_front_leg_tip_posY_left_middle_leg_tip |
| 116 | r_posY_left_front_leg_tip_posY_left_rear_leg_tip |
| 117 | r_posY_left_front_leg_tip_posY_right_front_leg_tip |
| 118 | r_posY_left_front_leg_tip_posY_right_middle_leg_tip |
| 119 | r_posY_left_front_leg_tip_posY_right_rear_leg_tip |
| 120 | r_posY_left_middle_leg_tip_posY_left_rear_leg_tip |
| 121 | r_posY_left_middle_leg_tip_posY_right_front_leg_tip |
| 122 | r_posY_left_middle_leg_tip_posY_right_middle_leg_tip |
| 123 | r_posY_left_middle_leg_tip_posY_right_rear_leg_tip |
| 124 | r_posY_left_rear_leg_tip_posY_right_front_leg_tip |
| 125 | r_posY_left_rear_leg_tip_posY_right_middle_leg_tip |
| 126 | r_posY_left_rear_leg_tip_posY_right_rear_leg_tip |
| 127 | r_posY_right_front_leg_tip_posY_right_middle_leg_tip |
| 128 | r_posY_right_front_leg_tip_posY_right_rear_leg_tip |
| 129 | r_posY_right_middle_leg_tip_posY_right_rear_leg_tip |
| 130 | r_speed_left_front_leg_tip_left_middle_leg_tip |
| 131 | r_speed_left_front_leg_tip_left_rear_leg_tip |
| 132 | r_speed_left_front_leg_tip_right_front_leg_tip |
| 133 | r_speed_left_front_leg_tip_right_middle_leg_tip |
| 134 | r_speed_left_front_leg_tip_right_rear_leg_tip |
| 135 | r_speed_left_middle_leg_tip_left_rear_leg_tip |
| 136 | r_speed_left_middle_leg_tip_right_front_leg_tip |
| 137 | r_speed_left_middle_leg_tip_right_middle_leg_tip |
| 138 | r_speed_left_middle_leg_tip_right_rear_leg_tip |
| 139 | r_speed_left_rear_leg_tip_right_front_leg_tip |
| 140 | r_speed_left_rear_leg_tip_right_middle_leg_tip |
| 141 | r_speed_left_rear_leg_tip_right_rear_leg_tip |
| 142 | r_speed_right_front_leg_tip_right_middle_leg_tip |
| 143 | r_speed_right_front_leg_tip_right_rear_leg_tip |
| 144 | r_speed_right_middle_leg_tip_right_rear_leg_tip |
| 145 | r_acc_left_front_leg_tip_left_middle_leg_tip |
| 146 | r_acc_left_front_leg_tip_left_rear_leg_tip |
| 147 | r_acc_left_front_leg_tip_right_front_leg_tip |
| 148 | r_acc_left_front_leg_tip_right_middle_leg_tip |
| 149 | r_acc_left_front_leg_tip_right_rear_leg_tip |
| 150 | r_acc_left_middle_leg_tip_left_rear_leg_tip |
| 151 | r_acc_left_middle_leg_tip_right_front_leg_tip |
| 152 | r_acc_left_middle_leg_tip_right_middle_leg_tip |
| 153 | r_acc_left_middle_leg_tip_right_rear_leg_tip |
| 154 | r_acc_left_rear_leg_tip_right_front_leg_tip |
| 155 | r_acc_left_rear_leg_tip_right_middle_leg_tip |
| 156 | r_acc_left_rear_leg_tip_right_rear_leg_tip |
| 157 | r_acc_right_front_leg_tip_right_middle_leg_tip |
| 158 | r_acc_right_front_leg_tip_right_rear_leg_tip |
| 159 | r_acc_right_middle_leg_tip_right_rear_leg_tip |
